# Increased receptor affinity and reduced recognition by specific antibodies contribute to immune escape of SARS-CoV-2 variant Omicron

**DOI:** 10.1101/2022.03.11.483934

**Authors:** Anne-Cathrine Vogt, Gilles Augusto, Byron Martina, Xinyue Chang, Gheyath Nasrallah, Daniel E. Speiser, Monique Vogel, Martin F. Bachmann, Mona O. Mohsen

**Affiliations:** Department of BioMedical Research, University of Bern, Bern, Switzerland; Department of Immunology RIA, University Hospital Bern, Bern, Switzerland; The Jenner Institute, University of Oxford, Oxford, UK; Erasmus Medical Center, Department of Viroscience, Rotterdam, The Netherlands; Artemis Bio-Support, Delft, The Netherlands; Biomedical Research Complex, Qatar University, Doha, State of Qatar

**Author notes:** ^*^**Corresponding author**: Mona O. Mohsen.

**Keywords:** Omicron, Delta, SARS-CoV2, antibody

## Abstract

In this report, we mechanistically reveal how the Variant of Concern (VOC) SARS-CoV-2 Omicron (B.1.1.529) escapes neutralizing antibody responses, by characterization of this variant, and wildtype Wuhan and Delta variant (B.1.617.2). Convalescent sera as well as sera obtained from participants who received two or three doses of mRNA vaccines (Moderna-mRNA-1273^®^ or Pfizer-BNT162b2^®^) were used for comparison in this study. Our data demonstrate that both the Delta as well as Omicron variants exhibit higher affinity for the receptor ACE2, facilitating infection and causing antibody escape by receptor affinity (affinity escape), due to reduced ability of antibodies to compete with RBD-receptor interaction and virus neutralization. In contrast, only Omicron but not Delta variant escaped antibody recognition, most likely because only Omicron exhibit the mutation at position E484 associated with reduced recognition, resulting in further reduced neutralization (specificity escape). Nevertheless, the immunizations with RNA based vaccines resulted in marked viral neutralization *in vitro* for all strains, compatible with the fact that Omicron is still largely susceptible to vaccination-induced antibodies, despite affinity- and specificity escape.

## 1. Introduction

The appearance of the SARS-CoV-2 Omicron variant B.1.1.529 was reported in South Africa (SA) on 24^th^ November 2021 by the World Health Organization (WHO) (1). Two days later, Omicron was designated as a new variant of concern (VOC) due to its potential to spread rapidly worldwide. The first known sample was collected in SA on the 8^th^ November 2021 and the first sample detected outside SA was found in a traveler arriving in Hong Kong from SA via Qatar on 11^th^ November 2021. On 7^th^ January 2022, Omicron was confirmed in 135 different countries world-wide (2).

The WHO has estimated the danger from Omicron as potentially high due to the following reasons: the global risk of perpetuating SARS-CoV-2 pandemic remains high; recent data have indicated that Omicron shows a significantly higher R-value than the Delta VOC with already enhanced transmission (3, 4). Increased numbers of cases have previously led to increased hospitalization and mortality, threatening to overwhelm health-care systems. Preliminary evidence suggested an increased risk of reinfection with Omicron as compared to other VOCs (2). The concern was also growing because Omicron has also spread amongst doubly vaccinated people (5). In turn, COVID-19 caused by Omicron appeared relatively mild and health care systems have remained functioning despite huge numbers of infected individuals, making it possible that this new variant may help ending the pandemic by world-wide spreading and consequent broad immunization with significantly reduced disease severity and mortality.

Until recently, the Delta (B.1.617.2) VOC was the predominant SARS-CoV-2 variant before the emergence of Omicron. Delta VOC was discovered in late 2020 in India and had spread to more than 163 nations by 24^th^ August 2021. In June 2021, WHO has stated, that SARS-CoV-2 Delta strain would become the most prevalent strain in the world (6). Based on recent evidence, the Delta strain is 40-60% more transmissible than the previous forms such as wild type (WT) Wuhan or other recent VOCs such as the Alpha (B.1.1.7) VOC. Delta has also been associated with increased risk of hospitalization mostly for unvaccinated or partially vaccinated people (7) which appears different for Omicron. For clarification, the WHO has assigned the name Delta variant to lineage B.1.617.2 and it is one of the 3 Indian variants summarized under B.1.617. The Delta variant B.1.617.2 harbors seven mutations in the spike protein relative to the ancestor Wuhan SARS-CoV-2 strain; two of these mutations (L452R and T478K) are located in the receptor binding domain (RBD) region (6, 8). Importantly, the Delta VOC does not exhibit a mutation at position E484, which is usually associated with strong reduction of recognition by antibodies (9–11).

An unusually large number of 37 mutations has been identified in Omicron in comparison to other VOCs. Most mutations are located in the spike protein (5). Furthermore, 15 mutations have been detected in the RBD region which interacts with the main receptor angiotensin-converting enzyme 2 (ACE2) in comparison to only two mutations in the Delta variant (12). The spike protein, particularly RBD, is the main COVID-19 vaccine target (13, 14). Computational *in silico* modeling raised concerns about a possible ability of Omicron to dodge antibody-mediated immunity (15). Several reasons may account for the reduced ability of antibodies to neutralize emerging VOCs, including specificity alteration of antibody recognition (classical antibody specificity escape) as well as increased RBD-ACE2 affinity that partially outcompetes antibody binding (affinity escape) (10, 11, 16, 17). So far it seems clear that alteration of antibody recognition of Omicron is related to its E484A mutation (11, 17–19).

According to WHO, the global impact of Omicron will largely depend on four main aspects: (1) the transmissibility of the variant, (2) how well vaccinated and infected people are protected against Omicron, (3) the virulence of Omicron and finally (4) the awareness of the population to take protective measures (2). In light of the importance of these key parameters, our study aimed at revealing the mechanisms underlying the overserved increased transmissibility and reduced protection by vaccine- or infection-induced antibodies. Our data demonstrate an increased affinity of Omicron for the it`s receptor ACE2 and reduced recognition by serum antibodies. These factors cause reduced viral neutralization. From a mechanistical point of view, our data demonstrate that Omicron avoids viral neutralization by both classical change of antibody specificity, and affinity escape due to the high RBD-ACE2 affinity that outcompetes antibody binding (20).

## 2. Methods

### 2.1. Human sera

Human sera were collected from COVID-19 convalescent patients, vaccinated individuals who received 2 doses of mRNA vaccine (Moderna-mRNA-1273^®^ or Pfizer-BNT162b2^®^) and vaccinated individuals who received a total of 3 doses of mRNA vaccine (Moderna-mRNA-1273^®^ or Pfizer-BNT162b2^®^). Participants were recruited at University Hospital of Bern, Bern, Switzerland. Sera were collected within one-month post vaccination.

### 2.2. RBD-ACE2 binding kinetics

The binding kinetic was performed using Octet RED96E (Sartorius). SAX sensors were loaded with biotinylated ACE2 25μg/ml and subsequently quenched with biocytin. RBD was serially diluted in kinetics buffer and dissociation was performed in 300s.Kinetic buffer (KB) was used to dilute proteins. A loaded sensor run in KBserved as a control. The resulting curves were aligned to beginning of association, and a 1:1 model was used for global fitting.

### 2.3. RBD proteins

Recombinant SARS-CoV-2 spike S1 B.1.1.529-Omicron RBD protein was purchased from (antibodies-online GmbH), recombinant SARS-CoV-2 Spike RBD and recombinant SARS-CoV-2 Spike RBD B.1.617.2 L452R T478K were purchased from R&D systems. Proteins were reconstituted as per the manufacturer’s instructions.

### 2.4. Anti-RBD titers

ELISA assays were performed as follow; 96 half-well plate were coated over night with 1 μg/ml of RBD proteins. Plates were blocked for 2h with 0.15% casein in PBS and bound for 1h with 1:20 sera diluted in 1:3 steps. Bound IgG antibodies were detected using goat anti-human IgG-POX antibody (Nordic MUbio). ELISA plates were developed using tetramethylbenzidine (TMB) and stopped with 1 M H2SO4. Absorbance was read at 450nm and curves were generated using OD_450_ and OD_50_.

### 2.5. BLI-based competitive assay

The ability of the human sera to compete with ACE2 for binding to RBD_WT_, RBD_Delta_ and RBD_Omicron_ was tested as previously described by Vogel et al. (11).

### 2.6. Neutralization assay (Cytopathic Effect-Based Neutralization Assay)

Sera samples were heat-inactivated for 30min at 56°C and diluted from 1:20 to 1:320. 100 TCID50 of WT (SARS-CoV-2/ABS/NL20), Delta (SARS-CoV-2/ABSD/NL21) and Omicron (SARS-CoV-2/hCoV-19/NH-RIVM-72291/2021) was added to each well and incubated for 1h at 37°C. Following incubation, the mixture was added on a monolayer of Vero cells and incubated for additional 4 days at 37°C. Afterwards, wells were inspected for the presence of cytopathic effect (CPE) and titers were expressed as the highest serum dilution which fully inhibits CPE formation (100% inhibition).

### 2.7. Statistics

GraphPad Prism 9.0 (GraphPad Software, Inc) was used to perform statistics. *Paired or Unpaired* Student’s *t*-test was performed to test the statistical significance between two groups (as indicated in the figure’s legend). Statistical significance is displayed as p ≤ .05 (*), p ≤ .01 (**), p ≤ .001 (***).

## 3. Results

### 3.1. Omicron VOC accumulated many more mutations than the Delta variant

The Omicron variant (B.1.1.529) harbors thirty seven mutations in the spike protein, half of which are located in the RBD region (21). In contrast, Delta VOC (B.1.617.2) has only two mutations in its RBD protein, namely L452R and T478K. L452R mutation has been shown to be common in several newly emerged variants of SARS-CoV-2 including Epsilon, Kappa, Lambda and Iota. Within the RBD region, L452R mutation locates in the receptor binding motif (RBM) in immediate proximity to ACE2 and has been associated with stronger receptor binding and cellular as well as immune escape (9, 22). The Delta VOC (B.1.617.2) lacks E484Q mutation present in another Indian VOC (Kappa B.1.617.1) but has a unqiue T478K substitution which is not found in other Indian variants (B.1.617.1 and B.1.617.3) (23). Interestingly, T478K mutation has also been detected in the Omicron VOC. Similar to L452R, T478K is in the RBD of the spike protein, also located within RBM of ACE2, making direct contact with the receptor (23). Figure 1 and Table 1 show the mutations in the RBD region of Omicron (B.1.1.529), Delta (B.1.617.2), and for comparison the ancestor Wuhan. The numerous mutations found in the RBD region of Omicron in comparison to Delta suggests that the newly emerged variant may be more immunologically resilient to neutralization by vaccine-induced antibodies.

**Table 1:** Mutations in RBD protein in Delta (B.1.617.2) and Omicron (B.1.1.529) VOC.

| RBD<sub>Delta</sub> (B.1.617.2) | RBD<sub>Omicron</sub> (B.1.1.529) |
| --- | --- |
| L452R and **T478K** | K417N, N440K, G446S, S477N, **T478K**, E484A, Q493R, G496S, Q498R, N501Y and Y505H |

Identical mutations sites in both Delta and Omicron variants are shown in magenta.

**Figure 1.**
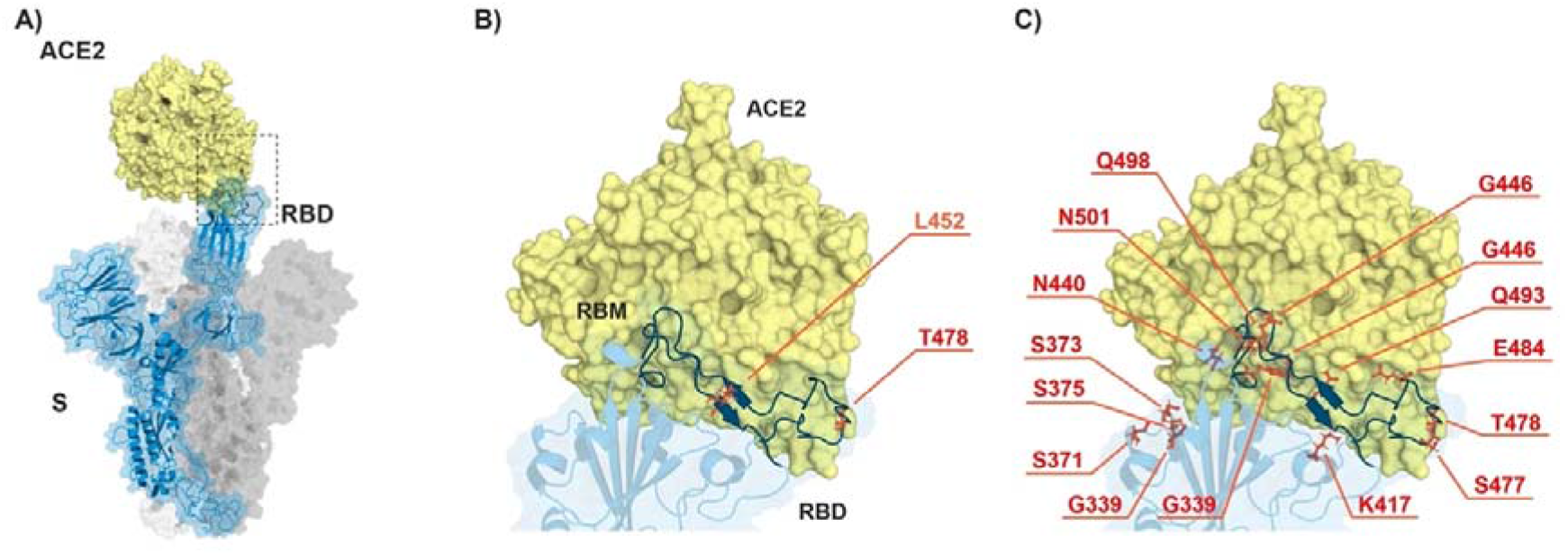
ACE2-spike interaction and mutations found in RBD of B.1.617.2 (Delta) and B.1.1.529 (Omicron). A) S monomer (blue ribbon and surface) bound to ACE2 ectodomain (yellow surface). B) Detail of A), highlighting the mutated residues (orange sticks) in Delta. C) Same as in B) but highlighting the mutated residues in Omicron. From PDB files 6ACG and 2AJF.

### 3.2. Binding kinetics of RBD Omicron (B.1.1.529) to ACE2 reveal similar affinities as RBD Delta (B.1.617.2) VOC

To address the binding properties of RBD_WT_, RBD_Delta_ and RBD_Omicron_ to the human receptor ACE2, we have utilized Biolayer Interferometry (BLI) to measure on- and off-rates as well as the affinity. The results confirm that RBD_WT_ has a binding affinity of K_D_ = 22.6 nM to ACE2 receptor (Fig. 2A) which is consistent with our previous findings (9, 11). Interestingly, RBD_Delta_ and RBD_Omicron_ exhibit an approximately similar two-fold increase in the binding affinity to ACE2 (RBD_Delta_ K_D_ = 10.5 nM, Figure 2B) and (RBD_Omicron_ K_D_ =11.6 nM, Figure 2C). The affinity increase of both RBD_Delta_ and RBD_Omicron_ is caused by an enhanced association rate and decreased dissociation rate (Table 2). These data indicate that RBD_Omicron_ and RBD_Delta_ show increased affinity to ACE2 compared to RBD_WT_. Interestingly, we have previously measured the affinity of RBD of another Indian variant (B.1.1617.1) harboring the two mutations (L452R/E484Q) and found that it was increased another 2-fold higher compared to the Delta variant ((9) and Table 2). As both Delta and Omicron became world-dominant and thus outcompeted this B.1.1617.1 variant, receptor affinity is apparently just one but not the most important parameter that drives efficiency of viral spread.

**Table 2:** Kinetic parameters for the RBD-ACE2 interaction calculated by BLI.

| Analyte* | $K_D$ [M] | $k_{on}$ [$M^{-1}s^{-1}$] | $k_{off}$ [$s^{-1}$] |
| --- | --- | --- | --- |
| RBD<sub>WT</sub> | $22.6 \times 10^{-9}$ | $1.7 \times 10^5$ | $3.9 \times 10^{-3}$ |
| RBD<sub>Delta</sub> | $10.5 \times 10^{-9}$ | $3.1 \times 10^5$ | $2.3 \times 10^{-3}$ |
| RBD<sub>Omicron</sub> | $11.6 \times 10^{-9}$ | $2.1 \times 10^5$ | $2.5 \times 10^{-3}$ |
| RBD<sub>L452R/E484Q</sub> (9) | $4.6 \times 10^{-9}$ | $7.2 \times 10^5$ | $3.3 \times 10^{-3}$ |

**Figure 2.**
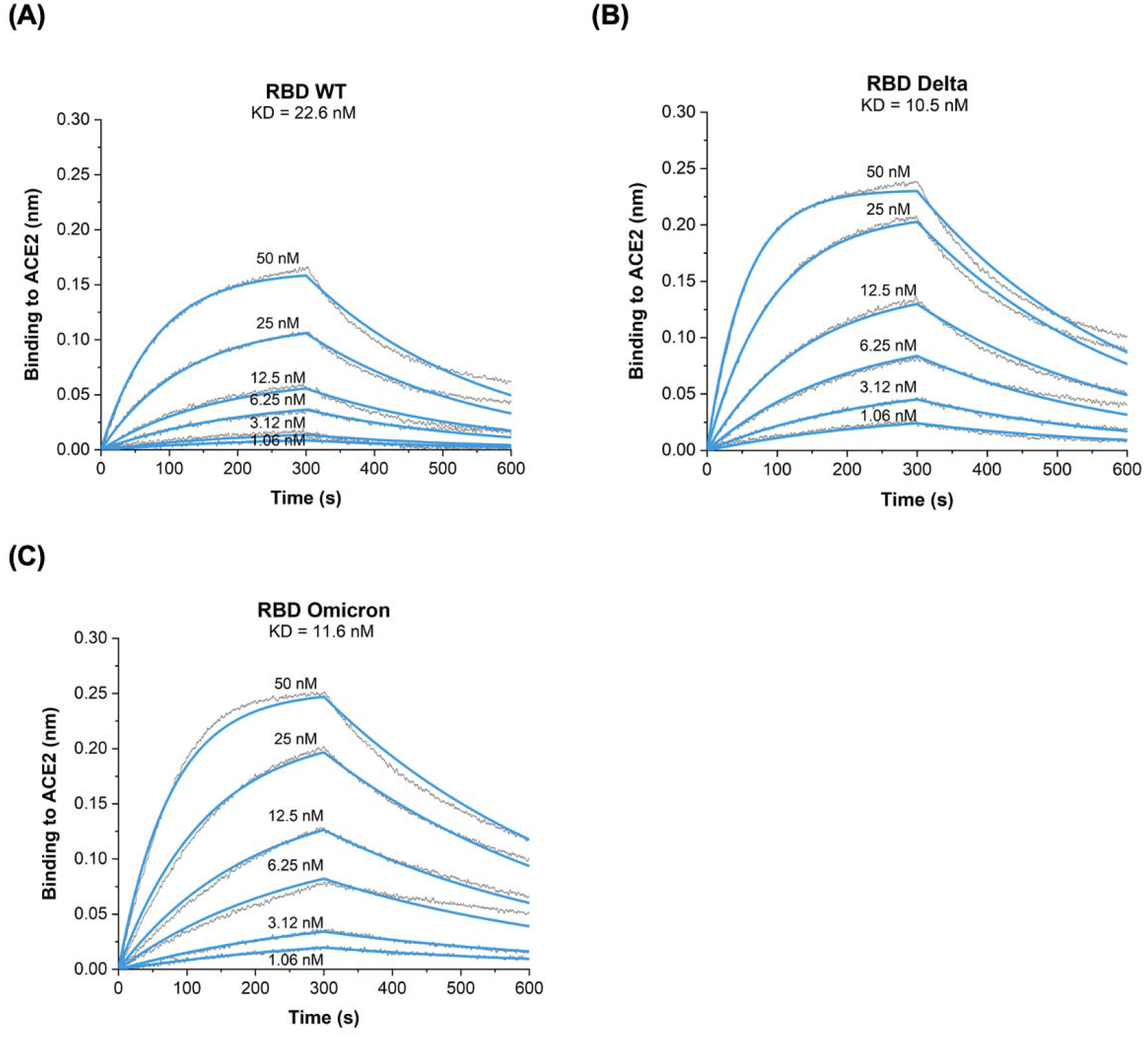
BLI sensograms illustrating ACE2 interactions with A) RBDWT; B) RBDDelta (B.1.617.2) and C) RBDOmicron (B.1.1.529). One representative of two similar experiments is shown.

### 3.3. Reduced recognition of Omicron mutant RBDs by sera from convalescent patients and mRNA vaccinated individuals

Here, we tested the ability of convalescent sera (Group I), sera from individuals after 2 doses of mRNA vaccine (Group II) and sera from individuals after 3 doses of mRNA vaccine (Group III) to recognize RBDDelta and RBDOmicron in comparison to RBDWT. Anti-RBD IgG ELISA revealed that antibodies induced by SARS-CoV-2 infection poorly recognize the different tested RBDs. On the other hand, Individuals who received 2 or 3 doses of mRNA vaccine could recognize RBDWT equally well to RBDDelta but recognition of RBDOmicron was reduced (Fig.3A-C).

When measuring the OD_50_ of the induced antibody titer, results indicate, as expected the superiority of the additional booster dose (Group III) (Fig. 3D). No statistical difference has been detected in recognizing RBDWT, RBDDelta or RBDOmicron after 2 doses of mRNA vaccine (Fig. 3E). However, a significant reduction in recognizing RBDDelta and RBDOmicron following 3 doses of vaccine (*p*. <0.0001) was stated (Fig. 3E). Analysis of AUC for convalescent demonstrates strongly impaired recognition of RBDOmicron but not RBD RBDDelta (Fig. 3F).

**Figure. 3.**
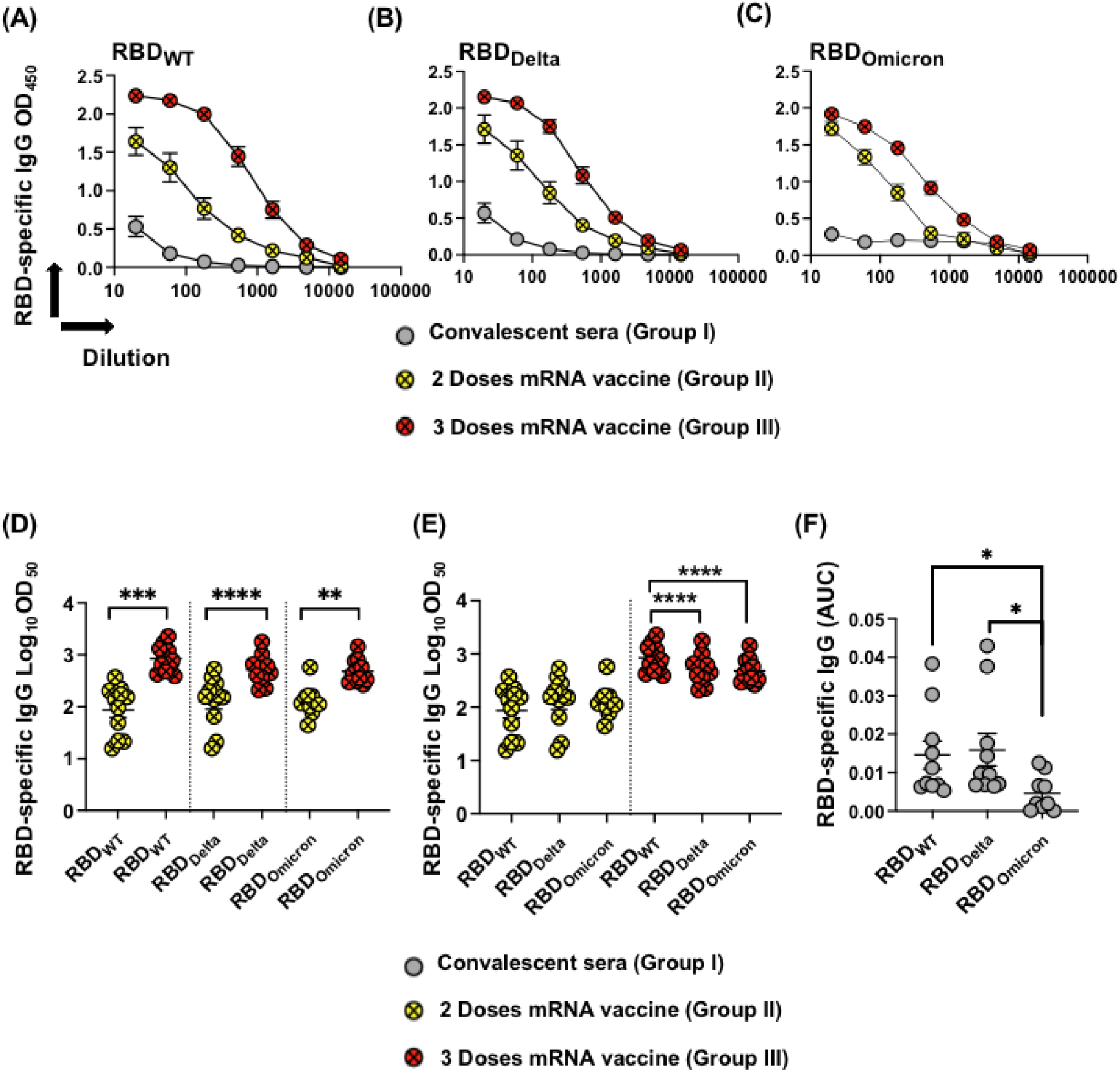
(A-C) Total IgG to RBD_WT_, RBD_Delta_ and RBD_Omicron_ of sera measured by OD_450_ obtained from COVID-19 convalescent donors (Group I), individuals after 2 doses of mRNA vaccine (Group II) and individuals after 3 doses of mRNA vaccine (Group III). (D and E) Log_10_ OD_50_ of total IgG to RBD_WT_, RBD_Delta_ and RBD_Omicron_, data from A-C (Groups II and III) measured by OD_50_. (F) AUC of IgG from convalescent patients to RBD_WT_, RBD_Delta_ and RBD_Omicron_, data from A-C (Group I). Statistical analysis by Student’s *t*-test, (Unpaired in D) and (Paired in E and F). N= 10 or 11. One representative of two similar experiments is shown.

### 3.4. Omicron RBD resists neutralization of receptor binding by immune sera

To further analyze RBD recognition by the different immune sera (Group I, II and III), we used Biolayer Interferometry (BLI) to measure antibody binding and the inhibition capability to block receptor interaction of the sera. As suggested by the ELISA results, there was a clear difference between recognition of RBDDelta and RBDOmicron, with the latter being recognized in an strongly inferior manner. Thus, recognition of the Omicron VOC is further reduced compared to Delta. As seen by ELISA, sera from individuals receiving 3 mRNA injections reacted best, flowed by those receiving 2 injections. The least reactive were convalescent sera (Fig. 4A and B).

**Figure. 4.**
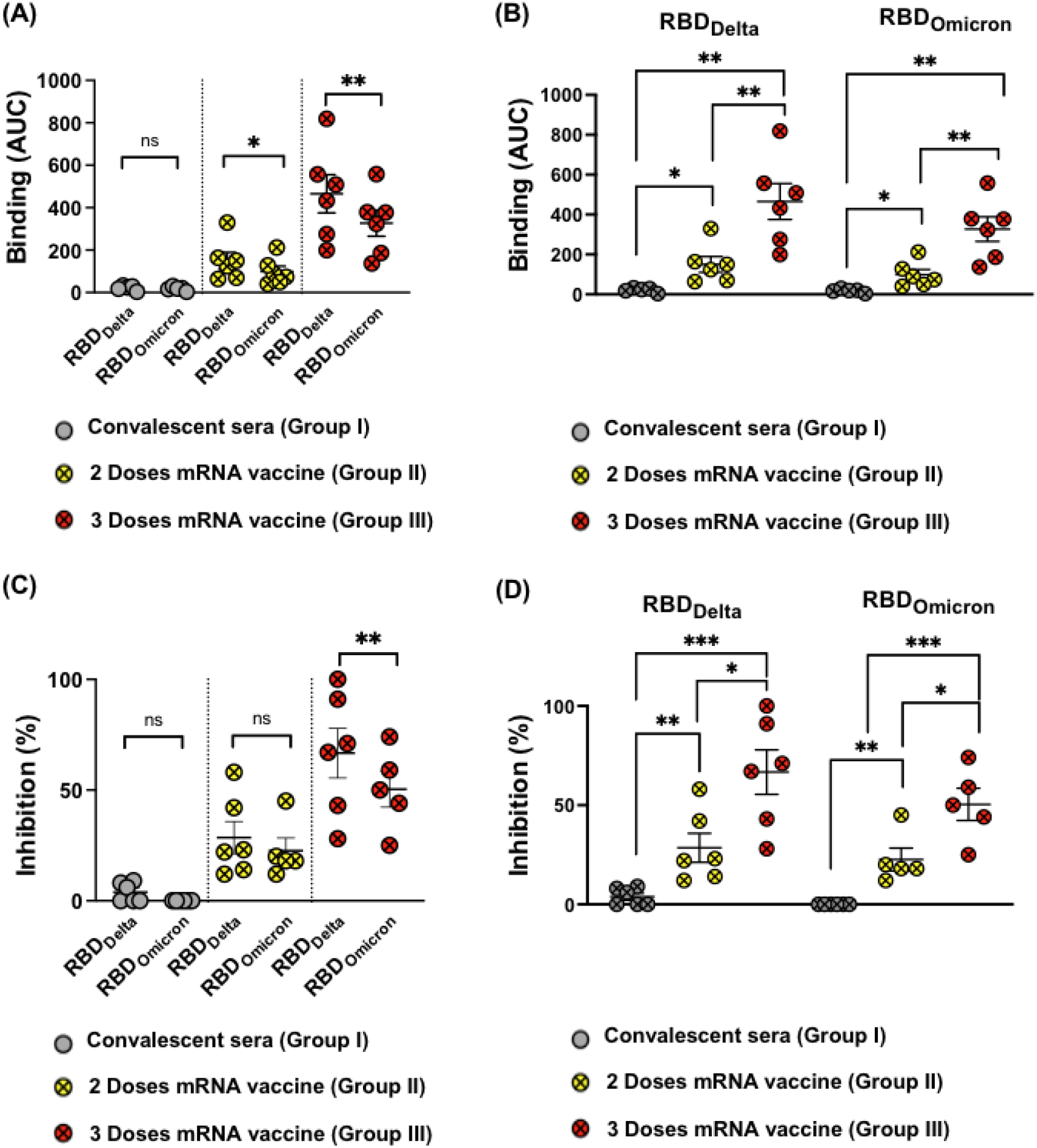
(A-B) Binding of sera from (Group I, II and III) to RBD_WT_, RBD_Delta_ and RBD_Omicron_ using BLI, depicted as AUC. (C and D) Inhibition of RBD_WT_, RBD_Delta_ and RBD_Omicron_ binding to the human receptor ACE2 by 1:80 sera dilution, sera from the three different groups were used in this assay. Statistical analysis by Student’s *t*-test, (Paired in A and C) and (Unpaired in B and D). N= 6 or 5. One representative of two similar experiments is shown.

Subsequently, we tested the ability of the different sera groups to inhibit RBDWT, RBDDelta and RBDOmicron binding to ACE2. Our results indicate that convalescent sera failed to effectively inhibit RBD-ACE2 binding at the concertation tested; such results are consistent with low induced antibody titers observed. Antibodies induced following the 2^nd^ dose of mRNA vaccination, show an average inhibition of 28.5% against RBDDelta versus 27.3% against RBDOmicron with no statistical difference (Fig. 4C). The overall average binding inhibition of the two VOCs to ACE2 receptor was dramatically increased after the 3^rd^ dose, accounting for 66% against RBDDelta but only 45% against RBDOmicron (*p*. 0.0011) (Fig. 4C). Thus, VOC Omicron evolved furthest to escape neutralization by antibodies induced by RBDWT. When comparing the inhibition data following 2 versus 3 doses of mRNA vaccine, the results revealed the significant ability of Group III in inhibiting RBDDelta and to a lesser extent RBDOmicron from binding to ACE2 receptor (Fig. 4D).

### 3.5. Omicron resists viral neutralization by immune sera

Cytopathic assay (CPE) was used to assess the neutralization capacity of the different sera Groups (I, II and III) against WT SARS-CoV-2 as well as the Delta and Omicron VOCs. Sera obtained from SARS-CoV-2 infected individuals who received no vaccination showed very low neutralization capacity against WT and both variants tested (Fig. 5A). Sera from individuals who received two doses of mRNA vaccine showed similar neutralization titers against both the WT and Delta with no statistical difference (*p*. 0.7961). In contrast to Delta, a significant reduction in the neutralization titer was observed against Omicron in comparison to WT (*p*. <0.0001) and Delta (*p*. 0.0011) (Fig. 5A). Such observation also applied to Group III for individuals who received 3 doses of the vaccine. A significant reduction in sera recognition to Omicron VOC was detected when compared to WT or Delta VOC (*p*. <0.0001).

**Figure. 5.**
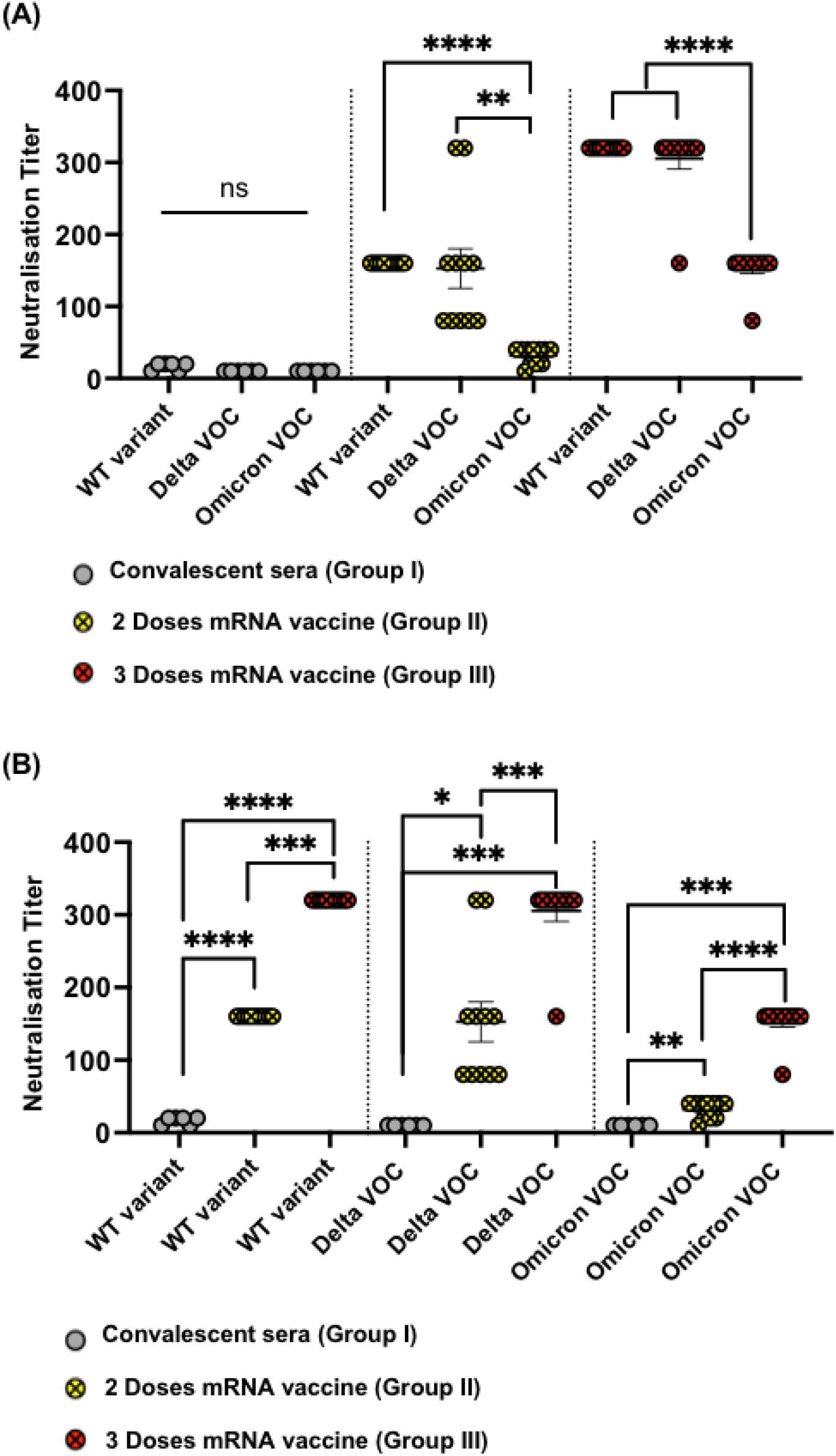
(A-B) Neutralization titers using CPE assay against WT, Delta and Omicron VOCs. Sera from Groups I, II and III were tested. Statistical analysis by Student’s *t*-test, (Paired in A) and (Unpaired in B). N=6 (Group I), N=11 (Groups II and III).

Next, we have analyzed the effect of the vaccination doses on each variant, i.e. a total of 2 doses vs 3 doses of mRNA vaccine. As expected, a booster dose of mRNA vaccine caused higher titers than natural infection with COVID-19. A second booster dose (i.e. 3 doses of mRNA) further enhanced the neutralization titer as shown in Figure 4B. The effect of a total of 2 or 3 doses of the vaccine was more variable and overall lower against Delta VOC than what was observed against the WT. Most importantly and as explained earlier, the overall neutralization titer against Omicron was reduced the most compared to the other variants (Fig. 5B).

### 4. Discussion

In this report, we have immunologically and mechanistically assessed the difference between the Delta VOC (B.1.617.2) with the Omicron (B.1.1.529) VOC using three different groups of sera. Omicron harbors 15 mutations in the RBD region beside other mutations in the spike protein. In contrast, Delta has only two critical mutations, L452R and T478K. The latter mutation has also been detected in Omicron. Both L452R and T478K mutations are located in the RBD, the essential region for viral binding to ACE2 and entry into receptor cells. The exact function of the remaining mutations in Omicron has yet to be revealed; nevertheless the unusually large number of mutations in Omicron raised serious concerns about reduced efficacy of the licensed COVID-19 vaccines to induce Omicron-neutralizing antibodies.

Recent studies have assessed the binding affinity of RBD_Omicron_ to human ACE2 receptor in comparison to the ancestral Wuhan strain, via atomistic molecular dynamics simulation (24) or surface plasmon resonance (5). The results revealed stronger binding to human ACE2 receptor, indicating that Omicron infects cells by a similar mechanism to the WT and suggesting that the clinically observed increased infectivity might be due to stronger binding interaction with ACE2. By performing Biolayer Interferometry (BLI) we demonstrate here definitively at the biophysical level that that both RBD_Omicron_ as well as RBD_Delta_ variant have a well-defined 2-fold increased affinity to ACE2 compared to RBD_wt_.

Our data show that Omicron has developed two mechanisms that account for the escape from neutralizing antibodies: First, some RBD mutations represent changes that result in antibody specificity alterations, such that antibodies generated by RBDs from previous infection or vaccination are less well blocking due to their specificity for previous RBD versions. Particularly the E484A mutation accounts for this type of specificity-escape, as it prominently changes an important epitope for neutralizing antibodies. Second, Omicron’s increased RBD-ACE2 affinity represents a challenge for neutralizing antibodies, as the virus’ RBD binds so strongly to ACE2 that antibodies become unable to compete and thus cannot block the virus. We call this phenomenon ‘affinity escape’ (19) as this type of immune escape is clearly different from the more classical antibody specificity escape.

Our findings are not surprising because Coronaviruses do not form serotypes, in contrast to Polio and Dengue viruses for example (25). Coronaviruses are large viruses with about three times more RNA nucleotides than most other RNA viruses. Despite the many small mutations that accumulate over time, Coronaviruses depend on a relatively high degree of genetic and structural stability which is assured by the RNA proofreading system of Coronaviruses (26). For enhancing transmission, an apparently better strategy than novel serotypes is to increase receptor affinity, a phenomenon that is now well documented for SARS-CoV-2 (9, 11, 27, 28). The new phenomenon of ‘affinity escape’ from neutralizing antibodies allows to better understand interactions of the virus with the immune system. We suggest that vaccines may be rendered more efficient by optimization for inducing large numbers of neutralizing antibodies with high affinity. It is well known that booster vaccinations significantly contribute to this aim, as continued affinity maturation in germinal centers of secondary lymphoid organs selectively generates and promotes high affinity antibodies (29). Indeed, individuals who have received three rather than only two RNA vaccinations reach relatively high protection from Omicron (30–32), although these vaccinations are still based on RBD_WT_ (Wuhan), supporting the notion that antibody specificity may not be the major reason for breakthrough infections.

In summary, both Delta and Omicron show enhanced receptor affinity translating into reduced neutralization (affinity escape). For Omicron, reduction in neutralization is, however, much more pronounced as its affinity escape is combined with specificity alteration of antibody recognition causing additional specificity escape.

## Acknowledgment

We thank Saiba GmbH for funding the project

## Authors contribution

Design of experiments, acquisition of data, interpretation and analysis of data: ACV, GA, BM, XC, MV, MFB and MOM. Writing, revision and editing of manuscript: MOM, DES, MV and MFB. FS and GN provided serum samples. Study supervision: MOM and MFB. All authors read and approved the final manuscript.

## References

1. Choudhary OP, Dhawan M, Priyanka. Omicron variant (B.1.1.529) of SARS-CoV-2: Threat assessment and plan of action. Int J Surg. 2022;97:106187.

2. Organization WH. Enhancing response to Omicron SARS-CoV-2 variant. 2022. p. https://www.who.int/publications/m/item/enhancing-readiness-for-omicron-(b.1..529)-technical-brief-and-priority-actions-for-member-states.

3. Gu H, Krishnan P, Ng DYM, Chang LDJ, Liu GYZ, Cheng SSM, et al. Probable Transmission of SARS-CoV-2 Omicron Variant in Quarantine Hotel, Hong Kong, China, November 2021. Emerg Infect Dis. 2021;28(2).

4. Clery D, Pennisi E. Omicron leads to fresh wave of meeting cancellations. Science. 2022;375(6577):132.

5. Mannar D, Saville Jw, Zhu X, Srivastava Ss, Berezuk AM, Tuttle KS, et al. SARS-CoV-2 Omicron variant: Antibody evasion and cryo-EM structure of spike protein-ACE2 complex. Science. 2022:eabn7760.

6. Mahase E. Delta variant: What is happening with transmission, hospital admissions, and restrictions? BMJ. 2021;373:n1513.

7. Bian L, Gao Q, Gao F, Wang Q, He Q, Wu X, et al. Impact of the Delta variant on vaccine efficacy and response strategies. Expert Rev Vaccines. 2021;20(10):1201–9.

8. Goher SS, Ali F, Amin M. The Delta Variant Mutations in the Receptor Binding Domain of SARS-CoV-2 Show Enhanced Electrostatic Interactions with the ACE2. Med Drug Discov. 2021:100114.

9. Augusto G, Mohsen MO, Zinkhan S, Liu X, Vogel M, Bachmann MF. In vitro data suggest that Indian delta variant B.1.617 of SARS-CoV-2 escapes neutralization by both receptor affinity and immune evasion. Allergy. 2022;77(1):111–7.

10. Mohsen MO, Balke I, Zinkhan S, Zeltina V, Liu X, Chang X, et al. A scalable and highly immunogenic virus-like particle-based vaccine against SARS-CoV-2. Allergy. 2022;77(1):243–57.

11. Vogel M, Augusto G, Chang X, Liu X, Speiser D, Mohsen MO, et al. Molecular definition of severe acute respiratory syndrome coronavirus 2 receptor-binding domain mutations: Receptor affinity versus neutralization of receptor interaction. Allergy. 2022;77(1):143–9.

12. Wu LY, Zhou LP, Mo MX, Liu TT, Wu CK, Gong CY, et al. SARS-CoV-2 Omicron RBD shows weaker binding affinity than the currently dominant Delta variant to human ACE2. Signal Transduct Tar. 2022;7(1).

13. Fantini J, Yahi N, Colson P, Chahinian H, La Scola B, Raoult D. The puzzling mutational landscape of the SARS-2-variant Omicron. J Med Virol. 2022.

14. Bachmann MF, Mohsen MO, Zha L, Vogel M, Speiser DE. SARS-CoV-2 structural features may explain limited neutralizing-antibody responses. NPJ Vaccines. 2021;6(1):2.

15. Nature. Heavily mutated Omicron variant puts scientists on alert. 2022. p. https://www.nature.com/articles/d41586-021-03552-w.

16. Liu X, Chang X, Rothen D, Derveni M, Krenger P, Roongta S, et al. AP205 VLPs Based on Dimerized Capsid Proteins Accommodate RBM Domain of SARS-CoV-2 and Serve as an Attractive Vaccine Candidate. Vaccines (Basel). 2021;9(4).

17. Zha L, Chang X, Zhao H, Mohsen MO, Hong L, Zhou Y, et al. Development of a Vaccine against SARS-CoV-2 Based on the Receptor-Binding Domain Displayed on Virus-Like Particles. Vaccines (Basel). 2021;9(4).

18. Chang X, Augusto GS, Liu X, Kundig TM, Vogel M, Mohsen MO, et al. BNT162b2 mRNA COVID-19 vaccine induces antibodies of broader cross-reactivity than natural infection, but recognition of mutant viruses is up to 10-fold reduced. Allergy. 2021;76(9):2895–998.

19. Bachmann MF MM, Speiser DE. To the editor: The Future of SARS-CoV-2 Vaccination. New England Journal of Medicine. 2022.

20. Bachmann MFM, M.O.; Speiser DE. The Future of SARS-CoV-2 Vaccination New England Journal of Medicinee. 2022.

21. Ren SY, Wang WB, Gao RD, Zhou AM. Omicron variant (B.1.1.529) of SARS-CoV-2: Mutation, infectivity, transmission, and vaccine resistance. World J Clin Cases. 2022;10(1):1–11.

22. Tchesnokova V, Kulakesara H, Larson L, Bowers V, Rechkina E, Kisiela D, et al. Acquisition of the L452R mutation in the ACE2-binding interface of Spike protein triggers recent massive expansion of SARS-Cov-2 variants. bioRxiv. 2021.

23. Di Giacomo S, Mercatelli D, Rakhimov A, Giorgi FM. Preliminary report on severe acute respiratory syndrome coronavirus 2 (SARS-CoV-2) Spike mutation T478K. J Med Virol. 2021;93(9):5638–43.

24. Lupala CS, Ye Y, Chen H, Su XD, Liu H. Mutations on RBD of SARS-CoV-2 Omicron variant result in stronger binding to human ACE2 receptor. Biochem Biophys Res Commun. 2022;590:34–41.

25. Plotkin SA. Updates on immunologic correlates of vaccine-induced protection. Vaccine. 2020;38(9):2250–7.

26. Sevajol M, Subissi L, Decroly E, Canard B, Imbert I. Insights into RNA synthesis, capping, and proofreading mechanisms of SARS-coronavirus. Virus Res. 2014;194:90–9.

27. Shang J, Ye G, Shi K, Wan Y, Luo C, Aihara H, et al. Structural basis of receptor recognition by SARS-CoV-2. Nature. 2020;581(7807):221–4.

28. Collier DA, De Marco A, Ferreira I, Meng B, Datir RP, Walls AC, et al. Sensitivity of SARS-CoV-2 B.1.1.7 to mRNA vaccine-elicited antibodies. Nature. 2021;593(7857):136–41.

29. Victora GD, Nussenzweig MC. Germinal centers. Annu Rev Immunol. 2012;30:429–57.

30. Garcia-Beltran WF, St Denis KJ, Hoelzemer A, Lam EC, Nitido AD, Sheehan ML, et al. mRNA-based COVID-19 vaccine boosters induce neutralizing immunity against SARS-CoV-2 Omicron variant. medRxiv. 2021.

31. Dejnirattisai W, Huo J, Zhou D, Zahradnik J, Supasa P, Liu C, et al. SARS-CoV-2 Omicron-B.1.1.529 leads to widespread escape from neutralizing antibody responses. Cell. 2022;185(3):467–84 e15.

32. Muik A, Lui BG, Wallisch AK, Bacher M, Muhl J, Reinholz J, et al. Neutralization of SARS-CoV-2 Omicron by BNT162b2 mRNA vaccine-elicited human sera. Science. 2022;375(6581):678–80.

